# scMoE: single-cell Multi-Modal Multi-Task Learning via Sparse Mixture-of-Experts

**DOI:** 10.1101/2024.11.12.623336

**Authors:** Sukwon Yun, Jie Peng, Namkyeong Lee, Yanyong Zhang, Chanyoung Park, Zunpeng Liu, Tianlong Chen

**Author notes:** Equal contribution (Sukwon Yun, Jie Peng). Correspondence to: Tianlong Chen < >.

## Abstract

Recent advances in measuring high-dimensional modalities, including protein levels and DNA accessibility, at the single-cell level have prompted the need for frameworks capable of handling multi-modal data while simultaneously addressing multiple tasks. Despite these advancements, much of the work in the single-cell domain remains limited, often focusing on either a single-modal or single-task perspective. A few recent studies have ventured into multimodal, multi-task learning, but we identified a ① **<u>Optimization Conflict</u>** issue, leading to suboptimal results when integrating additional modalities, which is undesirable. Furthermore, there is a ② **<u>Costly Interpretability</u>** challenge, as current approaches predominantly rely on costly post-hoc methods like SHAP. Motivated by these challenges, we introduce `scMoE`^1^, a novel framework that, for the first time, applies Sparse Mixture-of-Experts (SMoE) within the single-cell domain. This is achieved by incorporating an SMoE layer into a transformer block with a cross-attention module. Thanks to its design, `scMoE` inherently possesses mechanistic interpretability, a critical aspect for understanding underlying mechanisms when handling biological data. Furthermore, from a post-hoc perspective, we enhance interpretability by extending the concept of activation vectors (CAVs). Extensive experiments on simulated datasets, such as `Dyngen`, and real-world multi-modal single-cell datasets, including {`DBiT-seq, Patch-seq, ATAC-seq`}, demonstrate the effectiveness of `scMoE`. Source code of `scMoE` is available at: https://github.com/UNITES-Lab/scMoE.

## 1. Introduction

Given the inherently multi-modal nature of multi-omics, which includes transcriptome, genome, and proteome data at the single-cell level (Lee et al., 2020), there exists a notable mismatch with current methodologies. These methods are predominantly tailored for single-modality applications, targeting specific tasks (Van Dijk et al., 2018; Yun et al., 2023; Xiong et al., 2019; Cheung et al., 2021), thereby limiting their generalizability in a multi-modal environment encompassing diverse tasks. Such tasks encompass the identification of joint groups, such as cell type across different modalities, and cross-modal prediction, where one modality is utilized to infer the expression of cells in another. Recently, UnitedNet (Tang et al., 2023) proposed a multi-task learning framework given its multi-modal nature, employing an encoder-common fuser-decoder framework based on a shared latent space. However, this approach encounters two fundamental limitations:

### ① Optimization Conflict

As illustrated in Figure 1 (a), despite UnitedNet’s capability in handling a diverse multimodal environment, its peak performance is achieved using a subset of modalities (specifically, pre-MRNA and mRNA), rather than all four modalities, which paradoxically show the worst performance among the variations. This counterintuitive outcome, given the amount of information involved, is undesirable and highlights a limitation in harnessing the full potential of multi-modal data in the single-cell domain. A deeper investigation, as depicted in Figure 1 (b), reveals that the core issue stems from the encoder-common fuser-decoder framework in multi-task settings, leading to an optimization conflict. This conflict arises because the fuser consolidates information from each modality into a shared parameter space responsible for handling different tasks.

**Figure 1.**
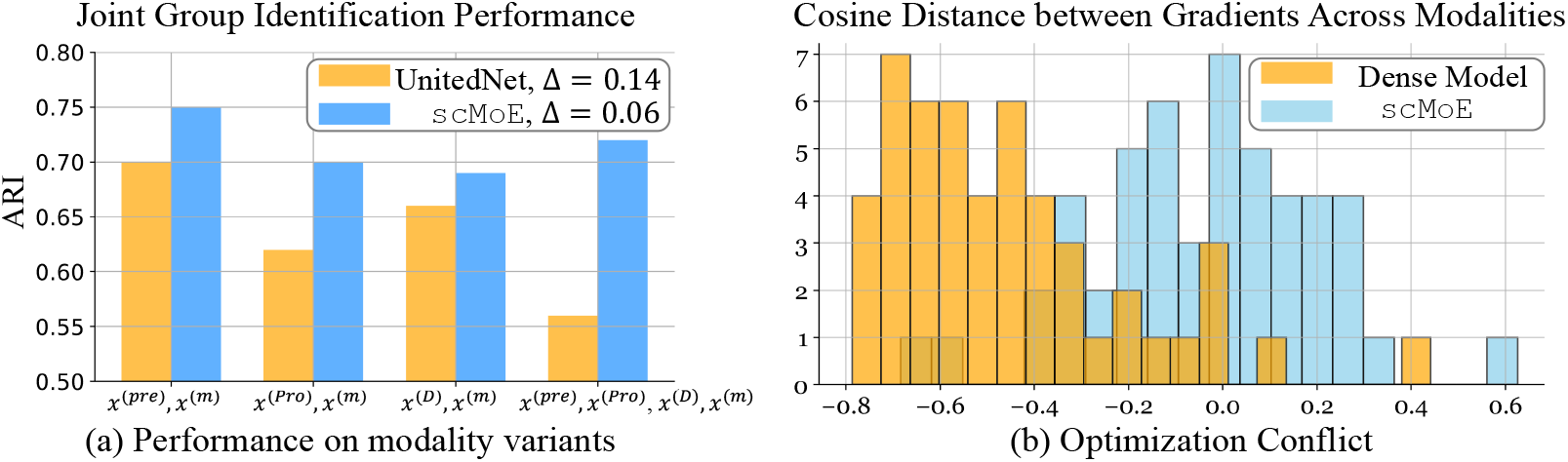
(a) Joint group identification performance across modality variants reveals that UnitedNet, when utilizing all modalities (Protein (*Pro*), mRNA (*m*), pre-mRNA (*pre*), DNA (*D*)), performs the worst. This is evidenced by a significant gap (Δ = 0.14) compared to our proposed model (Δ = 0.06), which maintains stable performance across a diverse range of modality combinations. (b) This phenomenon is attributed to optimization conflict issues, specifically gradient conflicts among modalities, such as “Protein” and “DNA” modalities. Here, the gradients are obtained from the experts and dense MLP with the same configuration in `scMoE` and Dense Model, respectively. Unlike currently adopted Dense Models like UnitedNet, our proposed sparse model demonstrates reduced conflict, as evidenced by more positive cosine distances, thereby facilitating enhanced multi-modal integration. The Dyngen dataset is used for the experiment.

This observation underscores the need for a framework designed to disentangle the parameter space, allowing for the coexistence of both common and specialized knowledge tailored for diverse tasks.

### ② Costly Interpretability

Interpretability in the biomedical and bioinformatics domain is essential, particularly for the practical application of these fields in clinical settings (Han & Liu, 2021; Karim et al., 2023). For instance, in predicting patient responses to cancer treatments using machine learning models, clinicians might hesitate to trust a model’s recommendations if they lack interpretability. A model that can elucidate the genetic markers or pathways influencing its predictions enables practitioners to make more informed, potentially life-saving decisions. Although UnitedNet provides some level of interpretability through post-hoc analysis with the SHapley Additive exPlanations (SHAP) algorithm (Lundberg & Lee, 2017), this method has its drawbacks. It comes with a high computational cost due to its need to evaluate all possible combinations of features, a complexity that increases exponentially with the number of features. This poses a significant challenge in the bio domain, where providing relevant explanations in a timely manner is crucial. This brings the necessity for mechanistic interpretability, which is inherently integrated into the model’s architecture, offering immediate insights during inference. Additionally, a more lightweight design for post-hoc analysis that aligns with the needs of the bio domain is also vital.

### ⋆ Sparse MoE as a solution

SMoE (Shazeer et al., 2017a) is an advanced neural network architecture that stands out for its ability to process complex, high-dimensional data efficiently. It achieves this by dynamically selecting a subset of specialized models, i.e., experts, for each input, offering a tailored approach in terms of a data-driven approach. Here, targeting challenges of single-cell multimodal data, we propose `scMoE`, which simply replaces the MLP layer in a transformer architecture with an SMoE layer so as to address the above limitations effectively. Specifically, it addresses the ① <u>Optimization Conflict</u> by employing multiple experts, thereby naturally disentangling the parameter space. This disentanglement allows for the attainment of both shared and unique knowledge tailored for each task and modality. Regarding the challenge of ① <u>Costly Interpretability</u>, `scMoE` addresses this by incorporating a gating network, or router, which automatically activates specific experts. This mechanism significantly enhances the model’s interpretability by immediately identifying which experts are most relevant for the task at hand during inference. Moreover, the integration of a cross-attention module before the SMoE layer, based on transformer architecture (Fedus et al., 2022), further enriches interpretability. This module adeptly captures the importance of feature combinations from different modalities, facilitating solving downstream tasks with improved efficiency and insights.

In summary, our contributions are three-fold:

- For the first time, we adopt the SMoE in single-cell multimodal multi-task learning to effectively tackle both optimization conflict and costly interpretability issues.
- To enhance interpretability efficiently, we investigate the use of concept-activation vectors (CAVs), which are particularly suitable for the single-cell domain.
- We demonstrate the effectiveness of `scMoE` across diverse multi-modal single-cell datasets. This includes simulations with the `Dyngen` dataset, as well as realworld datasets such as {`DBiT-seq, Patch-seq, ATAC-seq`}, in joint group identification and crossmodal prediction tasks.

## 2. Related Works

### Multi-Modal Multi-Task learning in single-cell data

Multi-modal learning (Makadia et al., 2008; Weston et al., 2011; Antol et al., 2015; Ramesh et al., 2022; Saharia et al., 2022; Yang et al., 2016; Dai et al., 2022; Jaegle et al., 2021) and multi-task learning (Xue et al., 2007; Hashimoto et al., 2017; Fan et al., 2022; Chen et al., 2023; Peng et al., 2024) have been subjects of extensive research over the years, with significant contributions from various fields. Such advancements inspired single-cell domain where uni-modal targeting signle-task, e.g., transcriptome with imputation task (Li & Li, 2018; Van Dijk et al., 2018; Wang et al., 2021; Yun et al., 2023) or clustering task (Tian et al., 2019; Lee et al., 2023), was predominant. For instance, MOFA (Argelaguet et al., 2020) disentangs variation in single-cell studies integrating different omics data types, like genomics and proteomics. totalVI (Gayoso et al., 2021), on the other hand, specifically integrates single-cell RNA sequencing data and protein abundance for a comprehensive cellular profile. WNN (Hao et al., 2021) combines single-cell RNA and protein data, creating a unified representation of cell states. Schema (Singh et al., 2021) integrates diverse singlecell omics data, including transcriptomics and electrophysiology, providing a holistic view of cellular function and state. Most recently, UnitedNet (Tang et al., 2023) has been introduced targeting multi-tasks like joint group identification and cross-modal prediction by utilizing a shared-latent space in a post-hoc explainable manner. However, as mentioned earlier, it encounters an optimization conflict issue. Involving more modalities can significantly degrade overall performance while also imposing a substantial burden on post-hoc interpretability.

### Sparse Mixture-of-Experts (SMoE)

SMoE model builds on the traditional Mixture-of-Experts (MoE) framework by introducing sparsity to enhance both computational efficiency and model scalability (Shazeer et al., 2017a; Chen et al., 1999). Unlike dense models, SMoE activates only a subset of relevant experts for each task, which reduces computational load and enables it to handle complex, high-dimensional data effectively. This selective activation has proven advantageous across diverse domains, including vision and language tasks, where it can dynamically adapt different parts of the network to specialized sub-tasks or data types (Riquelme et al., 2021; Lepikhin et al., 2021; Yun et al., 2024). While the model has shown success in traditional applications, its potential in bioinformatics and single-cell multimodal research remains largely untapped. Here, we explore SMoE’s capacity to model various modality combinations and address the challenges of missing modality scenarios within the biological domain.

## 3. Methodology

### 3.1 Preliminaries

#### Joint Group Identification with Cross-Modal Prediction

Given a multi-modal single-cell data, we aim to solve a multi-task problem. The first task, joint group identification, is to identify jointly expressed characteristics, a commonality across cells despite their differing modalities such as cell type, states, or tissue regions. From a classification perspective, both unsupervised and supervised approaches can be utilized simply by modifying the loss function. Simultaneously, we aim to address the cross-modal prediction task, which infers the information, i.e., expression of cells in one modality, using data from other modalities. Considering technical noise or properties that are difficult to measure, such predictions have the potential to significantly impact the real-world single-cell domain.

#### SMoE Notation

To address the optimization conflict issue identified previously, we implement SMoE to separate the parameter space across different tasks and modalities. In our model architecture, which is based on the transformer block, we substitute the traditional feedforward network with an SMoE layer, as depicted in Figure 2(b). Formally, the some comprises several experts, denoted as *f*_1_, *f*_2_, …, *f*_E_, where `E` represents the total number of experts, and a routing mechanism, ℛ, which selects experts in a sparse fashion. For a given embedding **x**, the top-*k* experts are engaged by ℛ based on the highest scores ℛ (**x**)_*i*_, where *i* indicates the expert index. This procedure is articulated as follows:

**Figure 2.**
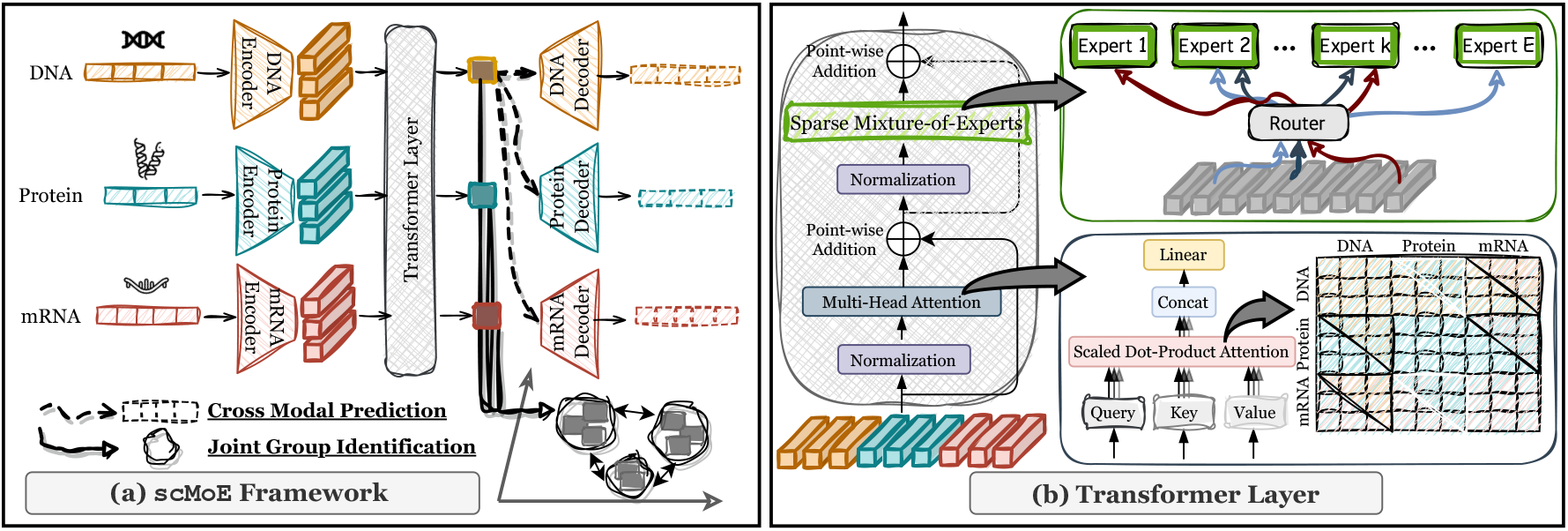
(a) The overview of `scMoE` : In the single-cell domain, each modality undergoes processing by its specific encoder, integrates with a shared transformer layer, and then passes through a modality-specific decoder. This structure enables simultaneous cross-modal prediction and group identification directly from the transformer output. (b) The transformer layer employs multi-head attention on concatenated modalities’ inputs, facilitating intra-modality self-attention and inter-modality cross-attention. A Sparse Mixture-of-Experts Layer then supersedes the standard feedforward network, enhancing the model’s efficacy in single-cell multi-modal multi-task learning.

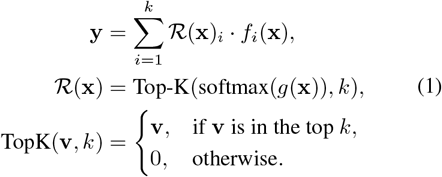

where **y**, the final output of the SMoE layer, is a weighted sum of the expert representations *f*_*i*_(**x**) and their corresponding weights ℛ(**x**)_*i*_, as determined by the router ℛ. Here, *g* denotes a trainable network, typically a small Feedforward Neural Network (FNN) that ranges from one to a few layers (Shazeer et al., 2017b; Riquelme et al., 2021). The `Top-K` (·) operation selectively retains a vector *v* if its probability is among the top *K* probabilities; otherwise, it sets the vector to zero.

### 3.2 scMoE : single-cell meets SMoE

In essence, as illustrated in Figure 2, `scMoE` adopts the **Encoder**-**Transformer**-**Decoder** framework. Below, we detail each module accordingly.

#### Encoder

Given the multimodal nature originating from diverse environments, such as multi-omics, we initially employ modality-specific encoders, denoted as *ε*^(1)^,…,*ε* ^(*V*)^, where 𝒱 represents the total number of modalities, to effectively generate embeddings informative of each modality. Notably, our use of transformer blocks (Fedus et al., 2022; Hu & Singh, 2021) requires input tokenization, differing from our matrix-formatted single-cell domain inputs where rows represent cells and columns denote specific modalities (e.g., genes or proteins), some of which, like highly variable genes (HVG) for the gene modality, vary in number. To achieve a consistent embedding shape across different modalities through tokenization, we adopt the patching method widely used in ViT-based works (Dosovitskiy et al., 2021). Thus, for an input **x**^(*ν*)^ ∈ ℝ^*B*×|*ν*|^, with batch size *B* and size of a specific modality |*ν*|, the output after processing by *ε*^(*ν*)^ is the embedding **h**^(*ν*)^ ∈ ℝ^*B*×*P*×*D*^, where *P* and *D* denote the desired number of patches and the hidden dimension, respectively. With these tokenized embeddings from each modality, we proceed to the transformer block, the core component of this work.

#### Transformer

The transformer block, depicted in Figure 2 (b), primarily functions as a feature extractor and includes two key components: (1) Multi-Head Attention, which facilitates both intra-modal and inter-modal attention through the modality-wise concatenated tokenized embeddings, **h** ∈ ℝ^*B*×*P*𝒱×*D*^. This setup enables the capturing of similarities between queries and keys across all modality combinations, with a total of 𝒱^2^. Representing the similarities in a matrix, diagonal elements would indicate selfattention within a modality (e.g., protein-protein) while offdiagonal elements signify cross-attention between different modalities (e.g., protein-mRNA), fostering a more thorough understanding of modalities. (2) The Sparse Mixture-of-Experts (SMoE) plays a crucial role in addressing optimization conflicts, thereby enhancing multi-task, multimodal environments as illustrated in Figure 1. By replacing the conventional Feedforward layer, SMoE enables the training of multi-experts who share common knowledge while retaining specialized knowledge in specific modalities or tasks. This is particularly pertinent in complex environments like the single-cell domain, where efficiency and specificity are essential. The output embedding from the transformer layer serves as a primary input for the unsupervised clustering loss, Deep Divergence-based Clustering (DDC) (Kampffmeyer et al., 2019), a method proven to enhance clustering in unsupervised settings. Notably, the DDC loss can be substituted with Cross-Entropy (CE) loss for supervised applications.

#### Decoder

In the single-cell domain, which often encounters noisy inputs due to dropout events (Hicks et al., 2018) and batch effects (Shaham et al., 2017), and lacks explicit supervision signals such as cell types, attaching decoder losses is a strategy to reconstruct the originally given input matrix effectively. Building on the final embeddings from the transformer block, we incorporate a total of decoders, 𝒟^(1)^, …, 𝒟^(*V*)^, mirroring the encoders. Given our focus on the cross-modal prediction task—predicting the expression of one modality (*ν*^2^) from another (*ν*^1^), where (*ν*^2^ ≠ *ν*^1^)—and considering that each modality serves as both input to itself and to other modalities, we aggregate a total of 𝒱^2^ reconstruction losses.

#### Training Procedure

Facing a multi-task learning scenario, we aggregate two primary losses: the DDC loss (or CE loss in supervised contexts) and the Reconstruction loss, addressing joint group identification and cross-modal prediction tasks, respectively. Unlike the iterative loss update strategy employed in UnitedNet (Tang et al., 2023), our method is straightforward, enhancing adaptability for future expansions to additional modalities. The comprehensive algorithm for training `scMoE` is detailed in Algorithm 1.

##### Algorithm 1

The overall procedure of `scMoE`

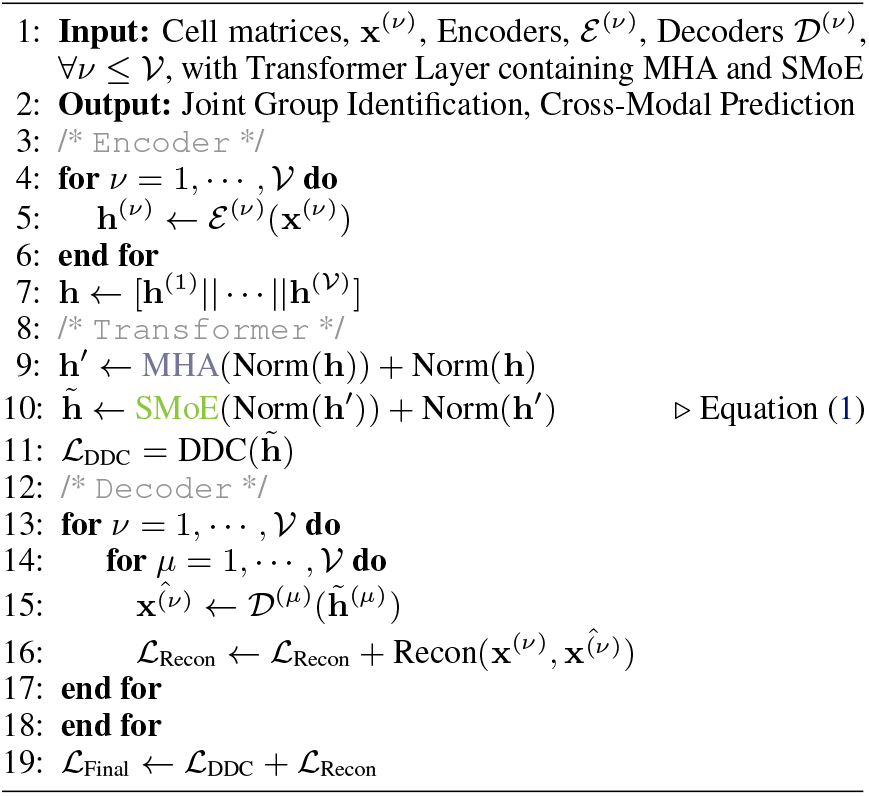

### 3.3 Interpretability of scMoE

In the field of biology, particularly in single-cell analysis, the interpretability of a proposed model is paramount. We explore the interpretability of `scMoE` from two perspectives: Mechanistic and Post-hoc.

#### Mechanistic Interpretability

Mechanistic interpretability (Wang et al., 2022; Kästner & Crook, 2023) involves understanding a model’s internal mechanisms and how they contribute to its decisions, observable during inference without additional training. While the integration of the SMoE layer might seem to obscure the model’s interpretability, the preceding multi-head attention mechanism, which captures both intra and inter-modality significance, maintains a level of interpretability. Furthermore, the SMoE layer’s gating network, or router ^2^, which decides which experts to activate for a given modality or task, allows for mechanistic interpretability through its data-driven decision-making process. This aspect will be demonstrated in the subsequent experimental section.

#### Post-hoc Interpretability

The mechanistic approach, while valuable, may not always be immediately transparent, as decisions such as expert selection are based on learned patterns. To complement the model’s complex inner workings without delving into intricate details, the post-hoc approach (Zhang & Zhu, 2018; Zou et al., 2023) provides insights at the input level. While SHAP (Lundberg & Lee, 2017) has been recently applied in single-cell multimodal analysis, its complexity prompts us to propose a more lightweight, directly applicable interpretability method based on Concept Activation Vectors (CAV) (Kim et al., 2018), tailored for the single-cell domain. Further details will be presented in Experiment Section 4.5.

## 4. Experiments

### 4.1 Experimental Settings

#### Datasets

To evaluate our method, we conducted experiments on **four** different datasets, including one simulated dataset and three real-world datasets. The simulated dataset is the Dyngen dataset, which contains 500 cells, each with DNA, pre-mRNA, mRNA, and protein modalities comprising 100 dimensions of features. This dataset was generated using the *Dyngen* software (Cannoodt et al., 2021). For real-world data, we utilized the Patch-seq GABAergic neuron dataset (*i*.*e*., PatchSeq dataset), which provides morphological (M), electrophysiological (E), and transcriptomic (T) features from GABAergic interneurons in the mouse visual cortex (Gouwens et al., 2020). After applying the quality control procedures from previous research (Gala et al., 2021), 3395 neurons were available for E-T analysis and 448 for M-E-T analysis. The Multiome ATAC+gene expression BMMCs dataset (*i*.*e*., ATAC-seq dataset) combines gene expression and genome-wide DNA accessibility data from 10 donors across 4 tissue sites (Luecken et al., 2021). The DBiT-seq embryo dataset (DBiT-seq dataset) includes 936 spots, with data on mRNA expression, protein expression, and niche mRNA. For more details about these datasets please refer to Appendix A.

#### Compared Methods

To demonstrate the superior performance of `scMoE` on the particularly challenging joint group identification task, we compare it with 5 state-of-the-art (SOTA) multi-modal integration methods: UnitedNet (Tang et al., 2023), the weighted Nearest Neighbor (WNN) (Hao et al., 2021), Schema (Singh et al., 2021), Multi-Omic Factor Analysis (MOFA) (Argelaguet et al., 2020), and totalVI (Gayoso et al., 2021). Additionally, we introduce an “Identification only” baseline for a more exhaustive comparison, which focuses solely on the identification aspect of the task without the complexity of integrating multiple modalities. Subsequently, we benchmark the cross-modal prediction performance of `scMoE` against three carefully selected baselines: UnitedNet, WNN, and the aforementioned “Identification only” method. We used the hyperparameter setting proposed in their paper, and for our best setting, please refer to Appendix B.

### 4.2 Simulation Study

As shown in Table 1, when targeting the unsupervised joint group identification task, our proposed method, `scMoE`, achieves the best scores across various combinations of modalities. Using solely pre-mRNA, Protein, DNA, and mRNA is more beneficial than using two modalities, such as DNA and mRNA. This finding contrasts with the recently proposed UnitedNet, which tends to fall short when incorporating more modalities. As corroborated by the optimization conflict issue illustrated in Figure 1, `scMoE` benefits from a performance gain when adding more modalities. Furthermore, in Table 4.1, which focuses on the cross-modal prediction task, `scMoE` consistently outperforms the baselines, demonstrating its effectiveness and suitability for multimodal and multi-task learning.

**Table 1.** Joint group identification task measured by ARI in simulated Dyngen dataset upon modality combinations. The performance is averaged upon 5-fold cross-validation sets.

|  | Dyngen |  |  |  |
| --- | --- | --- | --- | --- |
|  | Modality Combinations |  |  |  |
|  | Pre, m | Pro, m | D, m | Pre, Pro, D, m |
| scMoE | **0.75** | **0.70** | **0.69** | **0.72** |
| UnitedNet | 0.70 | 0.62 | 0.66 | 0.56 |
| Identification only | 0.49 | 0.41 | 0.44 | 0.50 |
| WNN | 0.43 | 0.46 | 0.43 | 0.46 |
| Schema | 0.55 | 0.57 | 0.56 | 0.56 |
| MOFA | 0.02 | 0.05 | 0.66 | 0.05 |
| totalVI | 0.07 | 0.02 | 0.02 | - |

**Table 2.**
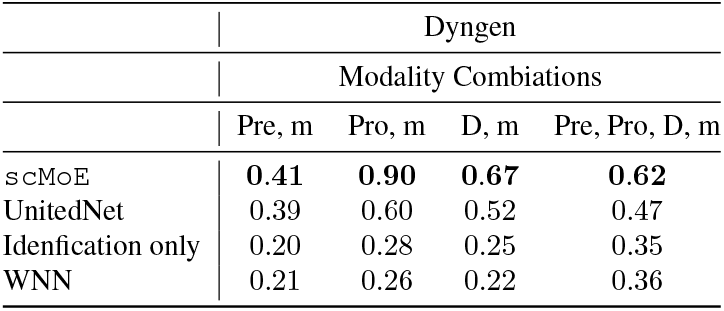
Cross-modal prediction task measured by *R*^2^ in simulated Dyngen dataset upon modality combinations. The performance is averaged upon 5-fold cross-validation sets.

**Table 3.** Ablation Study of scMoE.

|  | Dyngen |  |  |  |
| --- | --- | --- | --- | --- |
|  | Modality Combinations |  |  |  |
|  | Pre, m | Pro, m | D, m | Pre, Pro, D, m |
| scMoE | **0.97** | **0.83** | **0.89** | **0.75** |
| Dense | 0.68 | 0.68 | 0.71 | 0.72 |
| $N = 4$ | 0.82 | 0.71 | 0.77 | 0.75 |
| $N = 8$ | 0.80 | 0.69 | 0.76 | 0.75 |
| $N = 32$ | 0.74 | 0.65 | 0.76 | 0.74 |
| $k = 1$ | 0.79 | 0.68 | 0.73 | 0.70 |
| $k = 4$ | 0.82 | 0.70 | 0.79 | 0.74 |
| router per modality | 0.84 | 0.75 | 0.82 | 0.75 |
| 2 Patches per modality | 0.72 | 0.72 | 0.63 | 0.75 |
| 8 Patches per modality | 0.83 | 0.75 | 0.79 | 0.74 |
| 16 Patches per modality | 0.56 | 0.68 | 0.58 | 0.64 |

### 4.3 Real-world Applications

Figure 3 illustrates the performance of `scMoE` ĩn comparison with the recent SOTA, UnitedNet, across various datasets including DBiT-seq, ATAC-seq, and Patch-seq. Overall, `scMoE` ∼demonstrates significant generalizability across multiple multi-modal single-cell datasets, excelling in tasks such as unsupervised clustering and cross-modal prediction. Detailed settings and further insights are provided in the caption of Figure 3.

**Figure 3.**
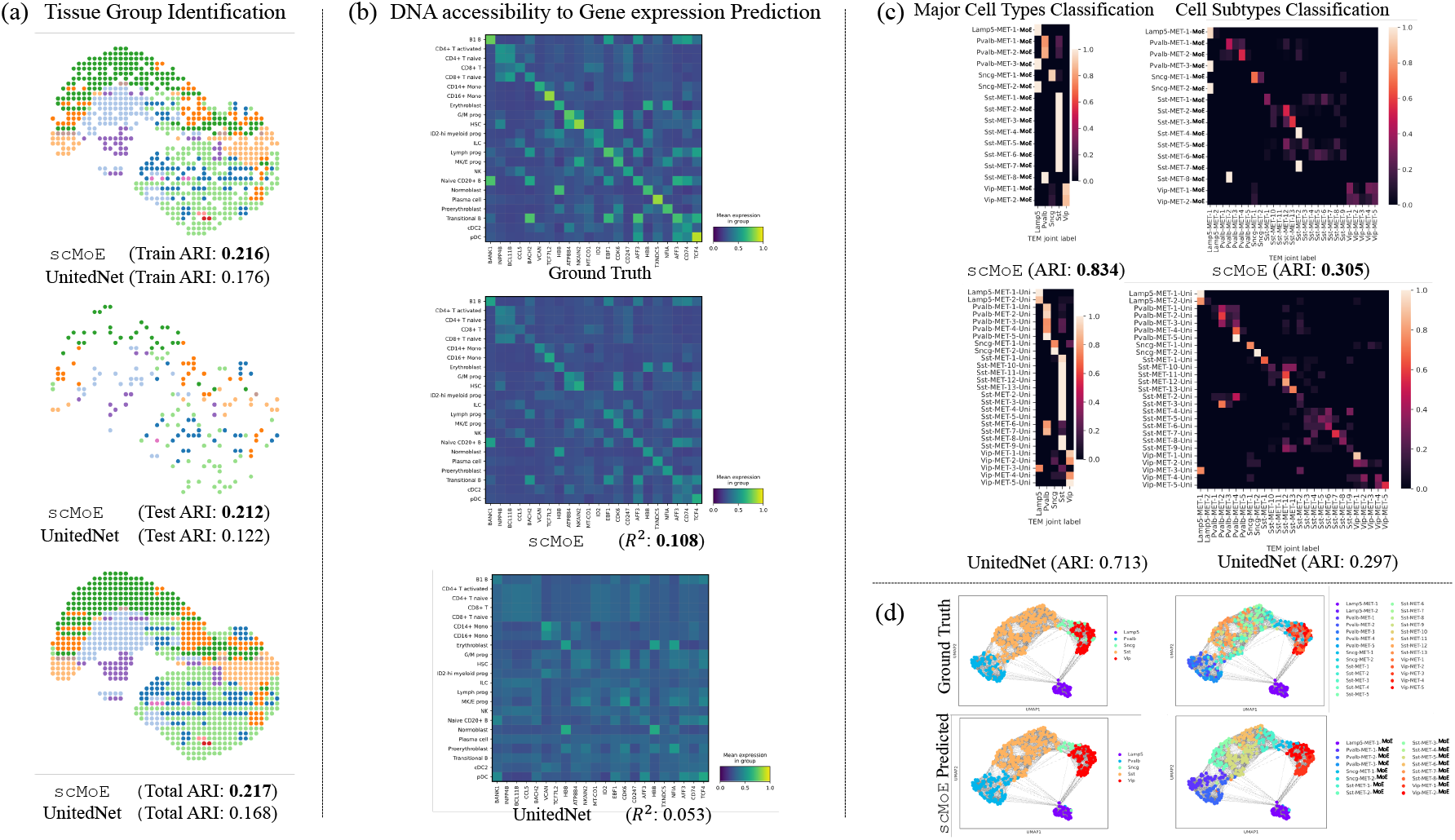
Performance analysis on real-world single-cell multimodal data: (a) From the DBiT-Seq dataset, we performed an unsupervised tissue group identification task to verify whether `scMoE` appropriately clusters 13 different groups. Compared to UnitedNet, our proposed model not only performs well in the training scenario but also in the unseen test scenario, verifying its generalizability. (b) Using the ATAC+gene expression BMMCs dataset, we conducted a supervised cross-modal prediction task to infer gene expression given DNA accessibility. Sharing a common trend in the representation of each cell type (row), thanks to the supervised signal, `scMoE` exhibits a similar gene expression trend in each cell type compared to the Ground Truth, outperforming UnitedNet. (c) Utilizing the Patch-seq dataset, we created a confusion matrix comparing the joint majority cell types and cell subtypes between reference labels and each model’s identified label (‘-MoE’ for `scMoE` and ‘-Uni’ for UnitedNet). Unlike UnitedNet, where uncertain predictions hamper overall performance, `scMoE` shows a notable performance gain, especially in major cell-type classification. This improvement is attributed to the gating network in the SMoE layer, making the decision process both efficient and effective. (d) Displays a UMAP representation of latent features colored by joint cell types, comparing the Ground Truth and `scMoE`.

### 4.4 In-depth analysis of Interpretability

#### Mechanistic Interpreatbility via SMoE and MHA

We detail our analysis on interpretability as illustrated in Figure 4. A principal aspect of mechanistic interpretability is its capacity to provide plausible reasoning without necessitating additional training. As depicted in Figure 4 (a), we emphasize that our experts within the SMoE framework can acquire both common knowledge (e.g., Expert indices 3 and 4 adept at handling Pre-mRNA and Protein modalities) and specialized knowledge tailored to each modality. This approach aligns with our aim to disentangle the parameter space to mitigate optimization conflict. Furthermore, as illustrated in Figure 4 (b), incorporating a Multi-Head Attention layer enhances mechanistic interpretability by underscoring the importance of considering inter-modal relationships in the multi-modal domain.

**Figure 4.**
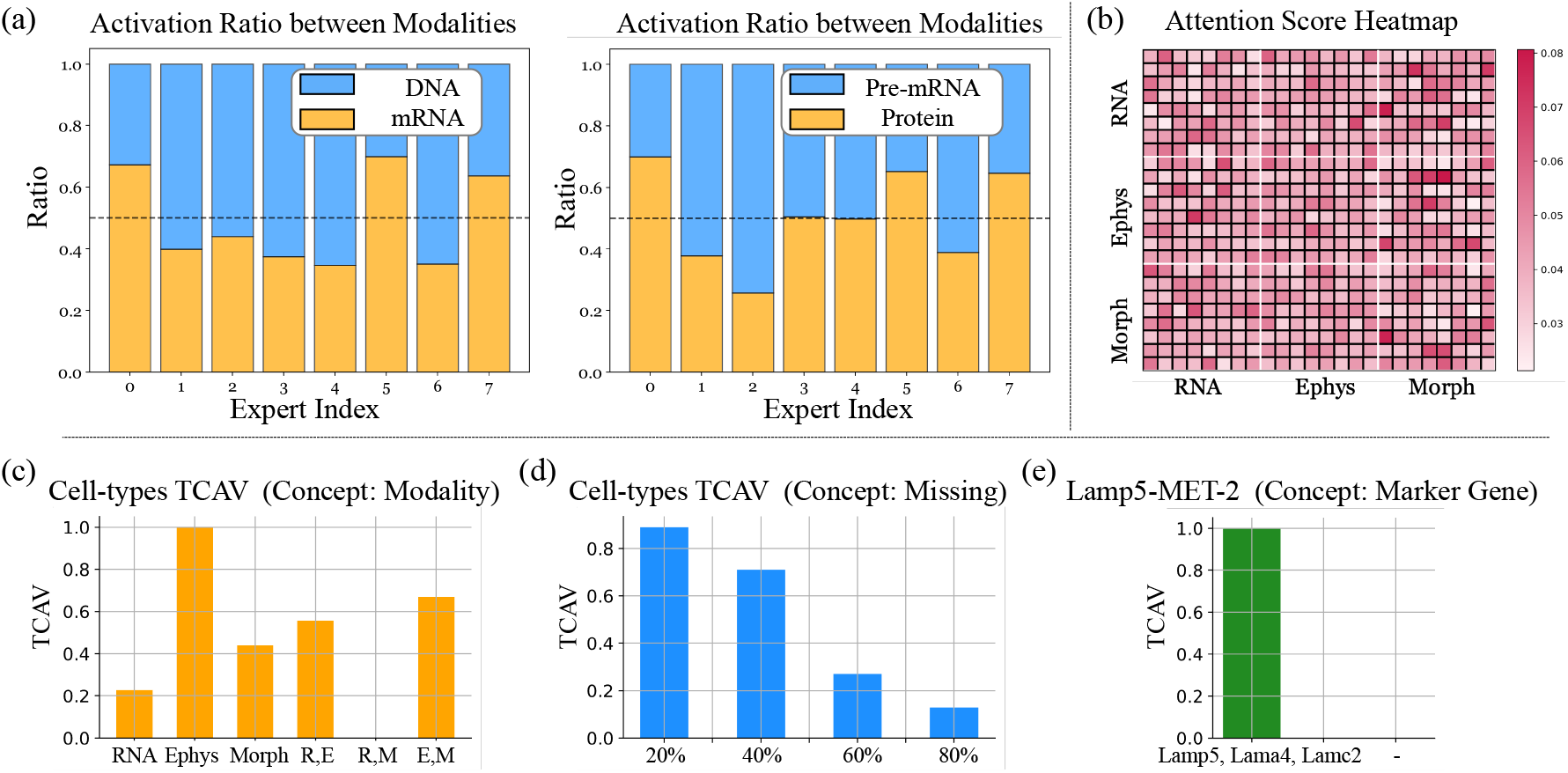
In-depth Analysis of Interpretability of `scMoE`. (a) Our framework, which adopts the SMoE approach comprising multiple experts and a gating network, dynamically utilizes the most beneficial expert for the given task at each iteration. We tracked this on-the-fly decision-making process and found that different experts specialize in different modalities. For instance, in the Dyngen dataset, experts 0, 5, and 7 showed heightened specialization in handling mRNA modalities, and vice versa for DNA. (b) An additional advantage of incorporating an attention module is observed through the consideration of both intra- and inter-modality relationships. Notably, significant cross-modality interactions, such as between Morph and Ephys, are evident in handling multi-modal data, as observed in the off-diagonal areas.(c) In terms of post-hoc interpretability, we introduce a novel approach utilizing TCAV, which offers new insights. Specifically, while classifying cell types by modality variants, we discovered that the Ephys modality plays a crucial role. (d) Additionally, in scenarios of RNA missing, commonly observed in the cell-gene matrix, we found that involving fewer missing nodes (20%) is advantageous for downstream tasks. Our TCAV analysis on identifying the rare cell type, i.e., ‘Lamp5-MET-2’, which constituted only 2 samples out of 448, involved defining the concept with widely recognized marker genes such as Lamp5, Lama4, Lamc2. Surprisingly, selective sampling of cells highly expressing these marker genes revealed that classifying this rare cell type is feasible despite the limited sample size, highlighting the significant role of marker genes in the single-cell domain. For (a), the Dyngen dataset was used, and for (b), (c), (d), and (e), the Patch-seq dataset was utilized.

#### Post-hoc Interpreatbility via TCAV

Next, to further enhance interpretability in the single-cell domain, we propose a novel approach based on Testing with Concept Activation Vectors (TCAV). More specifically, our goal is to improve model interpretability through the identification of high-level concepts using sets of example data.

To briefly introduce TCAV, a Concept Activation Vector (CAV) is generated by training a linear classifier to differentiate between examples of a concept and a random set of counterexamples. The CAV is then defined as the vector orthogonal to the classifier’s decision boundary. In the analysis of specific instances, TCAV calculates directional derivatives to assess how the model’s predictions are influenced by the core concept represented by the CAV. For instance, to evaluate the significance of the concept of stripes in an image of a zebra, a linear classifier is trained to distinguish between images of stripes and random samples. The CAV, orthogonal to the classification boundary, is utilized to gauge the sensitivity of the zebra image to the concept of stripes.

Inspired by TCAV, we propose a tailored approach for interpretability in the single-cell domain by designing concept vectors specifically suited for the biological domain. For example, leveraging the flexibility in defining concept vectors, we can identify which modality is crucial for a task or assess the impact of noisy and incomplete cell-gene matrix data, considering dropout events, or evaluate the efficacy of marker genes in classifying specific cell types, as illustrated in Figure 4. Furthermore, we wish to emphasize that the CAV approach is designed to be much more lightweight compared to other methods that rely on complex SHAP approaches, which consider all possible feature combinations. Instead, CAV provides a simpler mechanism by requiring only the training of a linear classifier based on a few subsets of specific samples of interest. Moreover, this CAV concept opens up numerous opportunities for new insights.

### 4.5 Ablation Study

To elucidate the contribution of the SMoE design within `scMoE`, we conduct a comprehensive series of ablation studies in Table 4.4 using the Dyngen dataset, specifically targeting the joint group identification task. We first examine the effects of SMoE by contrasting it with a dense model that possesses an equivalent number of parameters. Subsequently, our ablation analyses focus on the number of experts, the number of experts activated per token, the architecture of the routing network for SMoE, and the granularity of patches for each modality, aimed at understanding the impact on patch-level single-cell data representation. Within the Dyngen dataset, `SMoE` employs a single transformer block with the SMoE architecture. This layer incorporates 16 experts, with 2 activated experts per token. We configure the model to process 4 patches per modality and utilize a single routing network for expert selection. The SMoE architecture demonstrates superior performance compared to its dense counterpart indicating the SMoE mitigates gradient conflicts among diverse modalities. Additionally, our findings reveal that both an excessively large and an insufficiently small number of experts can harmfully impact the model’s performance. Similarly, the number of activated experts per token must be carefully calibrated, as too many or too few can also reduce model efficacy. We also observe that a modality-shared routing network outperforms a modality-specific routing network. Finally, our experiments identify that using 4 patches per modality represents a sweet point, with patch numbers beyond or below this threshold leading to diminished performance.

## 5. Conclusion

In this work, we investigate and design multi-modal multitask learning algorithms targeting high-dimensional singlecell data through the lens of Sparse MoE. Our pilot studies reveal two troublesome challenges in existing works, *i*.*e*., *optimizaiton conflict* and *costly interpretability*. To fill in the missing research gap, we tailor the sparse mixture-of-expert framework for single-cell data, which (1) disentangles the parameter space to allow modality- and task-specific modeling; (2) enjoys mechanistic interpretability inherently. Comprehensive validations on both simulated and real-world datasets consistently demonstrate the effectiveness of our proposal with enhanced interpretability. In the future, we are interested in extending our pipeline to systems immunology analysis with additional image modality.

## Impact Statements

The `scMoE` framework has the potential to advance personalized medicine by enabling precise single-cell data analysis, leading to more effective, patient-specific treatments and improved health outcomes, while supporting the broader goals of precision healthcare. Though we foresee significant societal benefits, we have not identified specific adverse consequences but remain committed to ongoing assessment of broader implications as the technology evolves.

## A Datasets and Preprocessing

### Dyngen Simulated Dataset

We use Dyngen (Cannoodt et al., 2021) to simulate a four-modality dataset comprising DNA, pre-mRNA, mRNA, and protein. The simulation generates 500 cells with each modality containing 100 dimensional features. Ground truth cell-type annotations are also provided. For Dyngen’s parameters, we adopt the default settings of a linear backbone model as outlined in the Dyngen tutorial, employing functions such as `backbone_linear, initialize_model`, and `generate_dataset`.

### Patch-seq GABAergic Neuron Dataset

We utilize a Patch-seq dataset from GABAergic interneurons in the mouse visual cortex. It contains 3395 neurons retained for E-T analysis and 448 for M-E-T analysis. We standardize the input matrices for each modality to normalize the mean and standard deviation of all features in each cell to 0 and 1, respectively.

### Multiome ATAC + Gene Expression BMMCs Dataset

We analyze a multiome ATAC + gene expression dataset from BMMC tissue across 10 donors and 4 tissue sites. Following quality control and standard preprocessing procedures, we normalize the gene expression data using median normalization, log1p transform, and standardization. We select the top 4000 most variable genes via Scanpy. For DNA accessibility data, we binarize the matrix and select the top 13, 634 most variable features, annotating DNA-accessibility peaks with ChIPseeker and `scanpy.var_names_make_unique`.

### UnitedNet on DBiT-seq Embryo Dataset

It includes three modalities: mRNA expression, protein expression, and niche mRNA expression from 936 spots. We normalize mRNA expression using `scanpy.pp.normalize_total` and select the top 568 differentially expressed genes. The protein expression is similarly normalized, focusing on 22 proteins, while niche modalities derive from normalized mRNA expression. For tissue region characterization, we extract ground truth labels from the original study and evaluate the clustering performance of UnitedNet against other methods using the adjusted rand index. For cross-modality prediction, we utilize mRNA and protein expression as inputs to UnitedNet. The DBiT-seq dataset is split into a training set (80%, 748 spots) and a testing set (20%, 188 spots) for the prediction task.

## B Hyper-Parameter Setting

### Dyngen

For the Dyngen dataset, we set the batch size to 64, the hidden dimension to 64, and train for 100 epochs. We represent each modality with 4 patches and use a learning rate of 1 × 10^−4^. The model’s architecture includes a single layer of transformer blocks with 1 attention head. In the Sparse Mixture-of-Experts (SMoE) component, we employ 16 experts, with 2 experts being activated simultaneously.

### DBiT-seq

For the DBiT-seq dataset, we configure the following parameters: a batch size of 64, a hidden dimension of 64, and 100 training epochs. Each modality is represented with 8 patches, and we employ a learning rate of 1 × 10^−4^. Our model architecture consists of a single layer of transformer blocks with 4 attention heads. In the Sparse Mixture-of-Experts (SMoE) component, we utilize 8 experts, with 2 experts being activated at a time.

### Patch-seq

The Patch-seq dataset experiments are conducted with a batch size of 32 and a hidden dimension of 128, over 100 training epochs. We represent each modality with 4 patches and set the learning rate to 4 × 10^−3^. The model includes one layer of transformer blocks with 4 attention heads. The SMoE component comprises 32 experts, with 2 experts activated simultaneously.

### ATAC-seq

In the ATAC-seq dataset, we use a batch size of 16 and a hidden dimension of 64, training for a total of 100 epochs. Modalities are represented with 8 patches each, and the learning rate is set to 1 × 10^−4^. The architecture includes a single layer of transformer blocks with 1 attention head. We employ 8 experts in the SMoE component, with 2 experts being activated per token.

## Footnotes

1 <u>s</u>ingle-<u>c</u>ell Sparse <u>M</u>ixture-<u>o</u>f-<u>E</u>xperts.

2 In this paper, we adhere to a single router shared across modalities to improve interpretability when analyzing experts.

